# Anti-Parkinsonian Drugs Rescue Locomotor Deficits in JIP3 Knockout Zebrafish: Implications for Treating Patients with *MAPK8IP3*-related Neurodevelopmental Disorders

**DOI:** 10.1101/2025.02.02.636112

**Authors:** Aleksandra Foksinska, Paige Souder, Gabrielle Smith, Kinnsley Travis, Sienna Rucka, Addie Allred, Rebeca Bender, Jackie Brunson, Avary Lanier, Amber Glaze, Andrew Crouse, Matt Might, Camerron M. Crowder

## Abstract

*MAPK8IP3-*related neurodevelopmental disorders are a spectrum of rare conditions caused by *de novo* mutations in the *MAPK8IP3* gene that encodes the JIP3 protein. These disorders are associated with a spectrum of neurodevelopmental symptoms that manifest in children and cause brain abnormalities, profound intellectual disabilities, movement disorders, and developmental delays. JIP3 is required for axonal transport of proteins and organelles between the soma and the synaptic terminal of neurons, a process critical for normal brain development and function. Homozygous loss-of-function mutations in JIP3 lead to impaired axonal transport and aggregation of cargo, which result in axonal swelling and stunted elongation. Despite these severe outcomes, disease mechanisms are poorly understood, and no current treatments are available.

Here we conduct thorough morphological, behavioral, and motility phenotyping in the JIP3 knockout zebrafish and identify locomotor deficits and morphological abnormalities. To identify treatment options, we used insights from expert clinicians and the artificial intelligence tool, mediKanren, to identify drug candidates hypothesized to improve patient symptoms or compensate for the loss of JIP3 at the molecular level. We then prioritized drugs that are FDA-approved, safe for children, and readily available. These collective efforts identified amantadine and levodopa as candidate therapies and rescued motor phenotypes associated with JIP3 loss-of-function in zebrafish.

## INTRODUCTION

Axonal transport is essential for proper neuronal growth and development. This process relies on motor proteins kinesin, dynein, and dynactin to transport cellular cargo along microtubules and scaffolding proteins to help dock and direct the cargo to the correct location within the axon. One of these scaffolding proteins, the JNK-interacting protein 3 (JIP3), encoded by the gene *MAPK8IP3* (mitogen-activated protein kinase 8 interacting protein 3), is highly expressed in the central nervous system and *de novo* variants are associated with a neurodevelopmental disorder, known as *MAPK8IP3*-related disorder (Iwasawa et al., 2019; Miura et al., 2006; Platzer et al., 2019).

### JIP3 structure and function

Key domains on the JIP3 protein allow for proper binding to various cargoes and appropriate motor proteins. The RH1 (RILP homology 1) and RH2 (RILP homology 2) domains are predicted to interact and bind to Rab GTPases and cytoskeletal motors such as myosin 5A(Vilela et al., 2019). Two leucine zipper domains, LZI and LZII, contain binding sites for kinesin heavy chain (KHC), kinesin light chain (KLC), dynactin, GTPase ARF6, and dynein light intermediate chain (DLIC) (Vilela et al., 2019). KHC and DLIC binding sites on JIP3 suggest critical roles in both anterograde (via kinesin-1) and retrograde (via dynein/dynactin) microtubule-based vesicular transport (Celestino et al., 2022). Accordingly, several studies report that JIP3 is essential for anterograde and retrograde transport of c-Jun N-terminal kinase (JNK), synaptic vesicles, tropomyosin receptor kinase B (TrkB), and endolysosomal organelles (Abe et al., 2009; Brown et al., 2009; Byrd et al., 2001; Celestino et al., 2022; Chen et al., 2015; Drerup & Nechiporuk, 2013, 2016; Edwards et al., 2015; Fu et al., 2022; Huang et al., 2011; Miller, 2017; Perlson et al., 2010; Platzer et al., 2019; Singh et al., 2024; Sun et al., 2017). Importantly, many cargoes associated with JIP3 are implicated in diverse neurodevelopmental and neurodegenerative disorders. Therefore, JIP3 protein dysregulation likely has broad negative impacts on neuronal health and function.

### JIP3 in neurodevelopment and neurodegeneration

JIP3 is posited to have crucial roles in both neurodevelopment and neurodegeneration. During development, kinesin-transported JIP3 activates JNK signaling at the axon tips, enhancing actin filament dynamics and axon elongation (Sun et al., 2013). Additionally, JIP3’s axonal transport of TrkB, a receptor for neurotrophins such as brain-derived neurotrophic factor (BDNF), contributes to the proper migration of cortical neurons (Ma et al., 2017). JIP3 also regulates neurite morphogenesis as a downstream effector of ARF6 and is essential for the distribution and number of axonal lysosomes, which support and maintain neuronal health and function (Ferguson, 2019; Suzuki et al., 2010). Additionally, JIP3 cargoes and interactors with established roles in neurodevelopment are also implicated in neurodegeneration. For instance, JNK signaling can be induced by cellular stress and initiate neuronal death while reduced BDNF levels or impaired BDNF/TrkB signaling is correlated with pre-clinical stages of Alzheimer’s disease (Ferrer et al., 1999; Kuan & Burke, 2005; Musi et al., 2021; Peng et al., 2005). Furthermore, loss of JIP3 in mouse neuron primary cultures and human induced pluripotent stem cells causes an accumulation of axonal lysosomes and elevated levels of Aβ peptide, derived from the Alzheimer’s disease-related amyloid precursor protein (Gowrishankar et al., 2017, 2021). Lastly, autophagic and endolysosomal pathways are implicated in neurodegenerative disorders such as Alzheimer’s and Parkinson’s disease, emphasizing the potential functional significance of JIP3 for maintaining neural integrity after development (Malik et al., 2019; Snead & Gowrishankar, 2022).

### *MAPK8IP3*-related disorder

Several *de novo* pathogenic variants in *MAPK8IP3* have been reported in children. These conditions are characterized by the National Organization for Rare Diseases (NORD) as neurodevelopmental disorders with or without variable brain abnormalities (NEDBA). Symptoms of these disorders include global developmental delay, hypo- and hypertonia, motor impairment, intellectual disability, poor or absent speech, brain and EEG abnormalities, short stature, facial dysmorphisms, early puberty, obesity, scoliosis, and cortical visual impairment (Iwasawa et al., 2019; Platzer et al., 2019). Most pathogenic *MAPK8IP3* variants appear to be missense variants, although a few are predicted to be null variants due to early truncations. Currently, disease-specific treatments are unavailable for this disorder. Additionally, how *de novo* variants yield neurodevelopmental or neurodegenerative phenotypes is unclear mechanistically. JIP3 KO mouse models (also known as JSAP1) displayed impaired axon projections, disorganized commissural tracts, and embryonic lethality, emphasizing the need for additional vertebrate models to characterize disease mechanisms. Furthermore, investigations into downstream pathways by observing how certain drugs impact disease models could lead to a rational basis for novel therapies.

In this research, we characterize motor and morphological phenotypes in JIP3 knockout zebrafish and attempted to rescue these phenotypes using AI- and physician-generated list of compounds. Here we present the outcomes of these drug treatments on locomotive activity in zebrafish and propose a hypothesis for ameliorating symptoms of *MAPK8IP3*-related disorder at the molecular level.

## MATERIALS AND METHODS

### Zebrafish Animal Husbandry

Zebrafish were housed in the University of Alabama Birmingham (UAB) Zebrafish Research Facility (ZRF). The ZRF uses a recirculating aquaria system (Aquaneering, Inc., San Diego, CA), and tank water is kept at 28°C. Zebrafish are maintained on a 14:10 light-dark cycle, fed 3 times daily, and observed by the UAB ZRF husbandry staff for any abnormalities. All methods were performed following guidelines provided by the UAB Institutional Animal Care and Use Committee.

All studies were conducted using a JIP3 knockout (KO) zebrafish line (JIP3^n17^), generated by Drerup *et al*. (2013) and provided to us by Dr. Nechiporuk from Oregon Health and Sciences University. These JIP3 KO fish were generated using ENU mutagenesis and found to have a single nucleotide substitution that resulted in an early stop codon that contained a SpeI restriction enzyme site. JIP3 heterozygotes adult fish were in-crossed approximately every 2 weeks to maintain the line. Heterozygous in-crosses were required due to breeding impairment of homozygous JIP3. Zebrafish embryos are kept in a 28.5°C Incubator in E3 water (603 E3B:17.2 g NaCl, 0.75 g KCl, 2.9 g CaCl2–H2O, 2.39 g MgSO4 dissolved in 1 L Milli-Q water; diluted to 13 in 9 L Milli-Q water plus 100 mL 0.02% methylene blue) until 5 days post fertilization (dpf). At 5 dpf, larvae are transferred to the recirculating water system in the ZRF nursery room.

### Genotyping

Genomic DNA was obtained by dissolving a whole embryo/larva or tail biopsies in 25-100 μL of embryo lysis solution (NaOH) from heterozygous in-crossed fish. Samples were incubated at 95°C for 25 minutes to extract genomic DNA. To neutralize the solution, 3 μL TRIS HCl solution was added and mixed thoroughly. Genotyping was conducted using standard polymerase chain reaction (PCR) performed as follows: 2x Taq master mix (New England Biosystems, Ipswich MA) with the following primers: forward – 5’-TTTGTCTGTTGAAATTGCT-3; reverse – 5’- ACGGTCCATACCCATGATT-3’, and genomic DNA following manufacturers protocol to generate at 385 base pair (bp) amplicon. PCR was performed using a Veriti™ 96- well thermal cycler from ThermoFisher Scientific and natural, semi-rimmed 96-well plates. The PCR reaction protocol was 95°C for 2 minutes, then 34 cycles of 95°C for 30 seconds, 55°C for 30 seconds, and 72°C for 1 minute and 30 seconds, and 72°C for 2 minutes. Following PCR amplification, an overnight (≥12 hour) restriction enzyme digest was conducted with SpeI/SpeI-HF following the manufacturer’s guidelines to generate double band homozygous amplicons (243 and 142 bp) easily detected using gel electrophoresis.

### Swim tunnel

Swim tunnel experiments were performed with the Loligo 170 mL swim tunnel system using Instacal, WitroxView, and AutoResp software. Adult (6-month-old) zebrafish were separated into breeding tanks (two fish per breeding tank with a clear barrier in-between) 24 hours prior to experiments for fasting. To minimize manipulation on experiment day, fish were measured and weighed during separation including weight, standard length, depth, and width. On experiment day, fish were transferred to the room with the swim tunnel in breeding tanks at least 1 hour prior to swim time for acclimation. The swim tunnel was filled with fresh system water maintained at ∼30°C and temperature was monitored throughout the experiment with a fully submerged thermometer. Measurements for each fish were entered into the software for critical swimming speed calculations prior to the experiment. To begin each swim experiment, fish were transferred individually into the swim tank apparatus and allowed to acclimate for at least 5 minutes prior to initiating flow. A modified aerobic closed respirometry protocol based on Marit and Weber 2012 was then performed, in which flow velocity was initiated at ∼2 body lengths (BL) per second for 5 minutes and increased by ∼1 BL/s in intervals of 5 minutes until exhaustion. Exhaustion was defined as the absence of swimming effort (fish resting against baffle) for a minimum of 5 consecutive seconds without renewed effort. Critical swimming speed (Ucrit) was calculated for each fish based on the equation in Plaut (2001).

### Behavioral Phenotyping

JIP3 heterozygotes were set up in breeding tanks in a 1:1 ratio the night before spawning in a single tank with a divider, to ensure breeding occurred in the morning. Heterozygous in crosses allowed for an unbiased screening of all three genotypes: wild type (JIP3 +/+), heterozygous (JIP3 -/+) and homozygous JIP3 knockouts (JIP3 -/-) clutch mates. Fertilized zygotes were collected within 2 hours of spawning time and transferred to Petri dishes (100 embryos per dish) with approximately 200 milliliters of E3 solution. Larval fish were incubated AT 28°C and E3 was replenished daily until 6 days post fertilization (dpf). At 6 dpf experimental assays were performed. Assays were repeated across genotypes, plates, and days and all assessments were completed during zebrafish light hours.

Locomotor phenotypes were first examined with the Noldus DanioVision system using the EthoVision XT v13 software (Basler GenICam Leesburg, VA). Larvae were allowed to acclimate in the DanioVision chamber for 10 minutes prior to the start of the assay. Larval swim patterns were recorded over a 1 hour and each well was inspected visually for 5 minutes to ensure successful tracking. Distance moved (mm), average velocity (mm/s), and cumulative time moving (s) were recorded based on tracking the center point of the individual larvae. Images with integrated visualization of the larval movement were also exported by the EthoVision XT tracking software. Images were captured at 1280×960 resolution and a sample rate of 25 samples per second. At 6 dpf, larvae were individually transferred to a 48-well round flat-bottom DanioVision plate using transfer pipettes. Wells were filled to 75% capacity with the E3 drug solution to similar levels.

The same larvae also underwent a second movement-based assessment with the Phylum Tech WMicrotracker ONE system and acquisition software, which allows for the assessment of 96 larvae at a time (Santa Fe, Argentina) (Bichara et al., 2014). This system has 2 microbeams per well and detects larva locomotor activity by counting interruptions or breaks in the light beams. Activity event parameters were defined as greater than 3% in the received light beam signal. Larvae were individually transferred to a 96-well flat-bottom plate using transfer pipettes. Wells were filled with E3 solution until they contained similar volumes. The larvae were allowed to acclimate in the room with the WMicroTracker ONE for 10 minutes prior to the assessment beginning. Events were measured in 30-minute intervals, over 2 hours, and averaged to calculate the average total locomotor activity. All assessments were completed during zebrafish light hours.

### Drug identification

Drug lists were generated using a variety of methods. Hypothesized JIP3 or related pathway modulators, drugs recommended to treat patient symptoms (i.e., rigidity, stiffness, spasticity, ataxia, ADHD), improve neuroplasticity, reduce inflammation or provide neuroprotection were queried and prioritized using the artificial intelligence tool, mediKanren (Foksinska et al., 2022). Pediatricians and neurologists treating *MAPK8IP3* patients also provided drug candidates for screening.

### Drug treatment

All drug compounds were ordered from Sigma-Aldrich or as libraries from MedChem Express. Drugs not soluble in E3 water were dissolved in a small amount of 0.05% DMSO and then brought up to stock concentration in E3 water solution. Additional lower concentrations tested were diluted from stock. Concentrations were selected from prior toxicology reports in zebrafish or tested through trial and error; most candidate drugs were screened at 0.1-100 µM. Drugs with less bioavailability or solubility were tested at higher concentrations. Zebrafish were incubated in petri dishes filled with drug E3 water immediately after fertilization and drug water changes were completed daily to replenish drug concentrations. Microtracker and DanioVision assays were performed at 6 dpf. Abnormal phenotypes or lethality were recorded.

### Statistical analysis

Analyses were conducted using GraphPad Prism software LLC (Boston, MA). Behavioral comparisons across genotypes were completed using 1-way ANOVAs with Tukey’s test for multiple comparisons. Swim tunnel groups (wild-type male, wild-type female, JIP3 -/- male, JIP3 -/- female) were compared using multiple unpaired t-tests, and significance was defined as a p-value less than 0.05.

## RESULTS

We first examined the impact of the loss of JIP3 on growth and morphological features throughout development we examined weight, standard length, depth, and critical swimming speed using the swim tunnel assay in male and female 6-month-old adult JIP3 KO zebrafish, compared to wild-type clutch mates (**Figure 3 A-F**). Weight and standard length, but not depth, were reduced significantly in adult males and females. Critical swimming was significant (**Figure 3 E**). Although slightly decreased in JIP3 KO males, only JIP3 KO females showed a significant decrease in critical swimming speed, compared to wild-type clutch mates (**Figure 3 F**). These findings indicate that JIP3 KOs display stunted growth and likely display sex-specific motility defects as adults.

**Figure 1.**
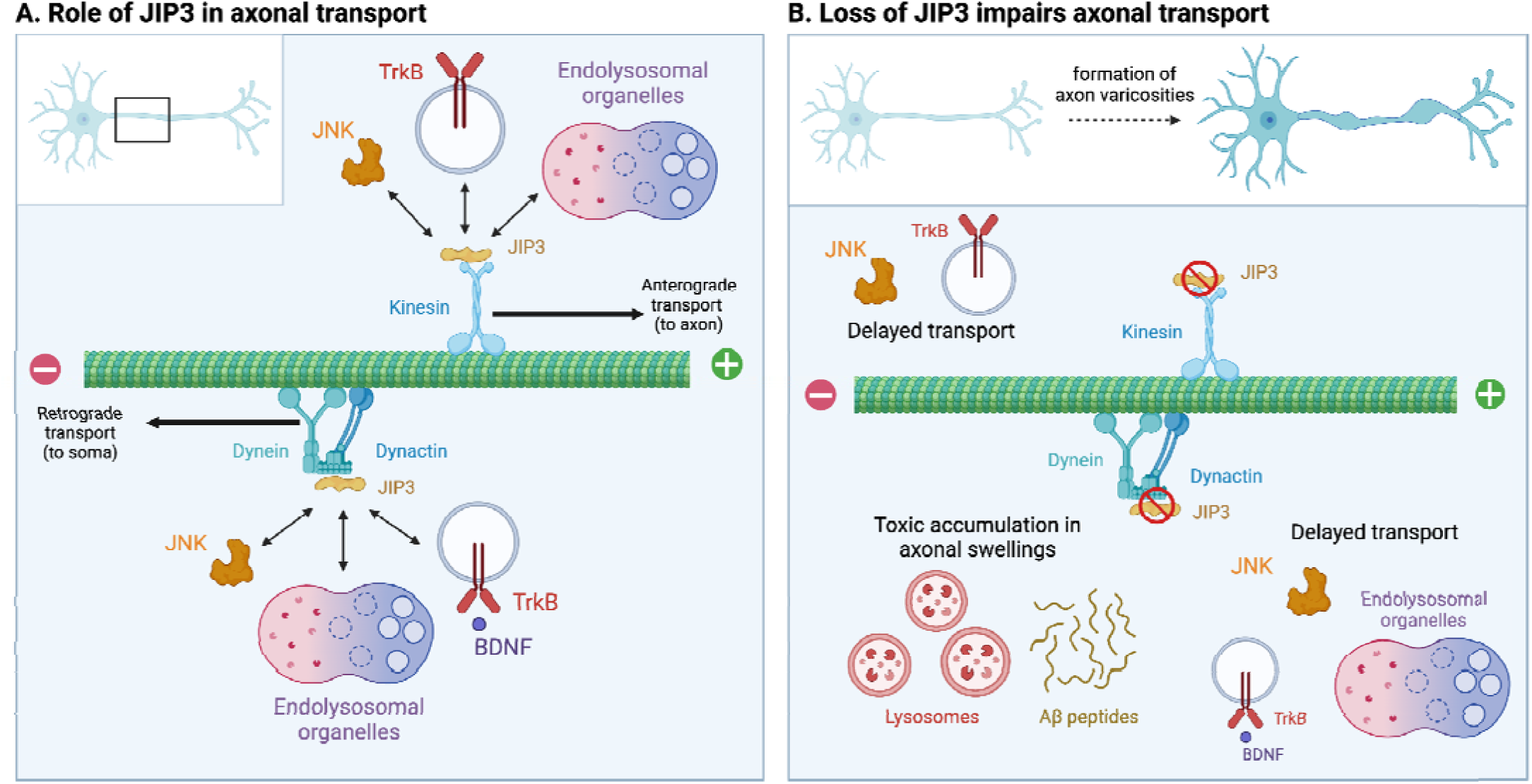
Knockout of JIP3 results in disrupted movement and accumulation of cargo in the axon. **A.** JIP3 acts as a scaffold protein binding to and aiding in the transportation of cargo, including other signaling molecules, such as JNK (Drerup & Nechiporuk, 2013; Sato et al., 2015; Sun et al., 2013) and TrkB (Huang et al., 2011), and endolysosomal organelles, such as autophagosomes (Cason & Holzbaur, 2023), lysosomes (Drerup & Nechiporuk, 2013), and early endosomes (Celestino et al., 2022). **B.** Loss of JIP3 function disrupts axon transport, leading to delayed transport and accumulation of lysosomes and amyloid precursor protein processing enzymes and Aβ peptides in axonal varicosities (swellings) (Gowrishankar et al., 2021). Created with Biorender.com.

**Figure 2.**
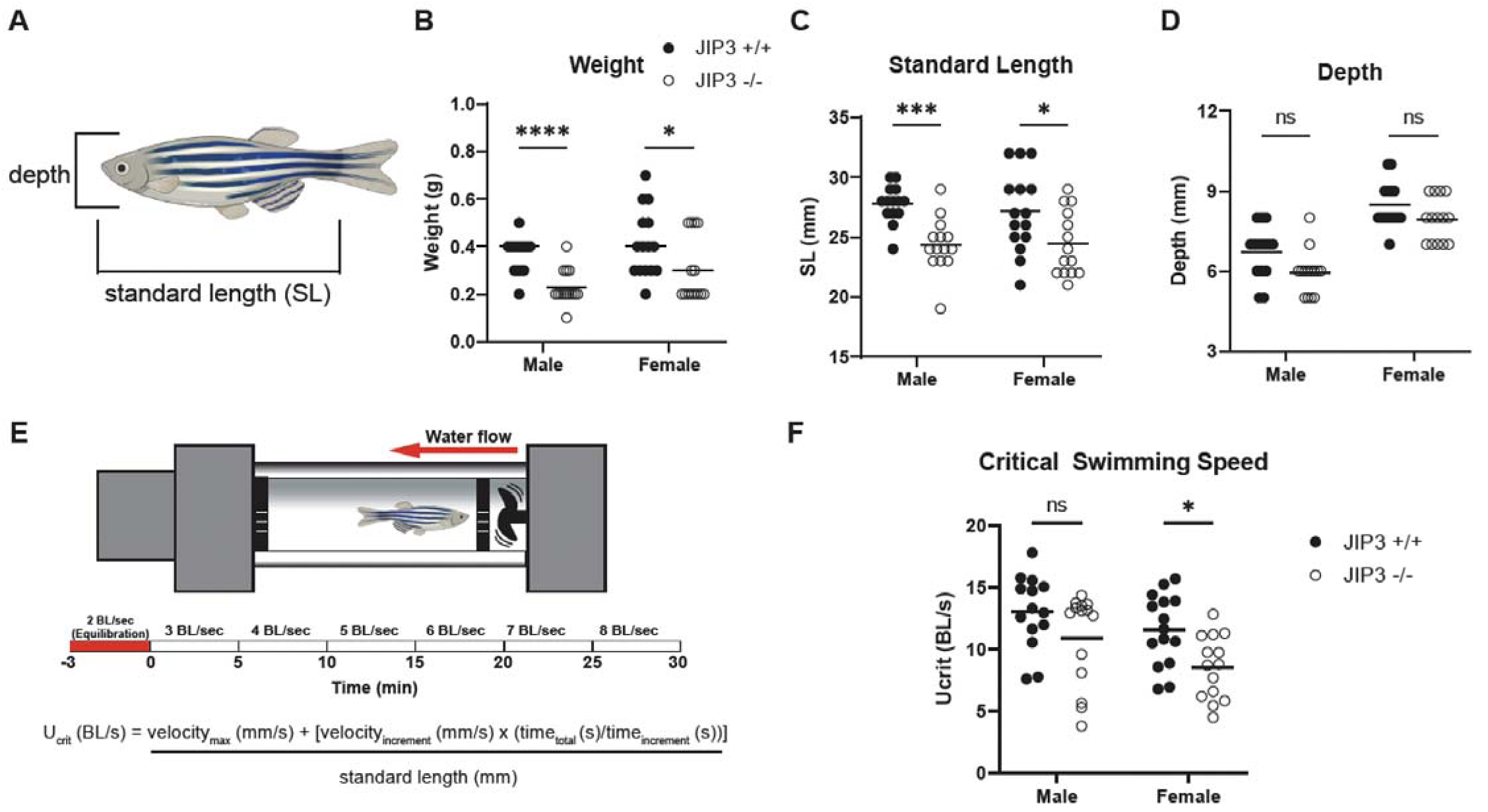
Adult weight, standard length, and critical swimming speed are decreased in JIP3 homozygous (-/-) adult zebrafish in a sex-specific manner. (**A**) Schema for standard length (SL) measures and depth for adult zebrafish. Measures from the tip of the snout to the base of the tail exclude the tail fin, and depth measures the widest point of the main body of the fish, excluding the pectoral fins. (**B-D**) Comparison of weight (**B**), standard length (**C**), and depth (**D**) of male and female wild type (JIP3 +/+) and homozygous JIP3 mutant (JIP3 -/-) 6-month-old adult zebrafish. Legend in (**B**) applies to **C-D**. (**E**) Schema for swim tunnel protocol to measure critical swimming speed in adult zebrafish. (**F**) Critical swimming speed in BL/s of male and female JIP3 +/+ and JIP3 -/- zebrafish revealed a significant decrease in females, but not males. Comparisons for all groups were made using multiple unpaired t-tests. *p<0.05, **p<0.01, ***p<0.001, ****p<0.0001, ns = not significant. Zebrafish images were created using Biorender.com.

**Figure 3.**
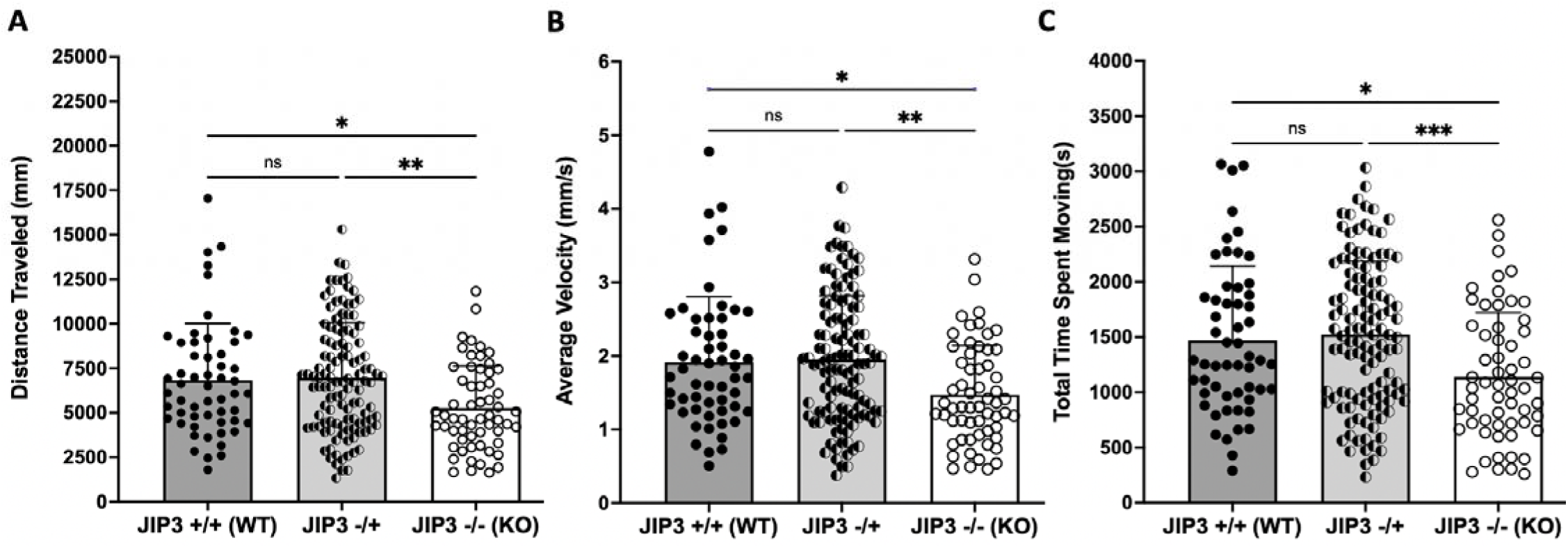
JIP3 KO larval zebrafish travel less distances, at slower velocities, and move for a shorter duration than wild type or heterozygous clutch mates. 1-hour DanioVision assay displaying decreased distance traveled (mm) (**A**), decreased average velocity (mm/s) (**B**), and decreased total time spent moving (s) (**C**) in JIP3 KO vs JIP3 heterozygous and wild-type clutch mates. Each dot represents an individual 6-day post-fertilization zebrafish larvae with mean and standard deviation provided. A 1-way ANOVA with Tukey’s test for multiple comparisons was conducted to compare means across genotypes, p-values are *p<0.05, **p<0.01, ***p<0.001, ns = non-significant.

To assess how JIP3 loss-of-function alters locomotor behaviors, we assayed 6-day post-fertilization larvae zebrafish using both the DanioVision and MicroTracker assays. DanioVision assays showed that JIP3 KO zebrafish traveled less distance (mm), at slower velocities (mm/s), and spent less time moving (s) than heterozygous or wild-type clutch mates (p < 0.05 - 0.001) (**Figure 4 A-C**). Likewise, higher throughput MicroTracker assays showed decreased average locomotor activity, compared to wild-type (p < 0.0001), and no difference was detected between JIP3 KO and heterozygous clutch mates (**Figure 5**). These data indicate that loss of JIP3 results in significant locomotor deficits in larval zebrafish.

**Figure 4.**
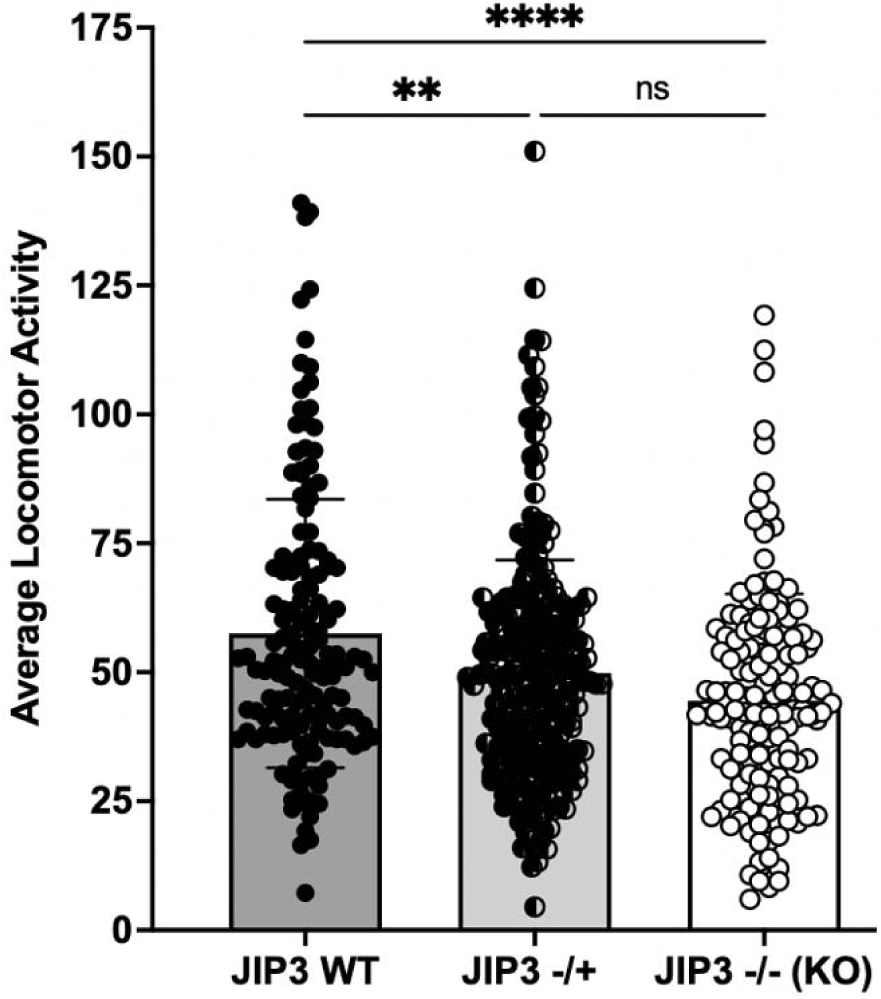
JIP3 KO larval zebrafish display decreased locomotor activity. A 2-hour MicroTracker 96-well plate assay displaying decreased average locomotor activity in JIP3 KO larvae, compared to heterozygous and wild-type clutch mates. Each dot represents an individual 6-day post-fertilization zebrafish larvae with mean and standard deviation provided. A 1-way ANOVA with Tukey’s test for multiple comparisons was conducted, p-values are *p<0.05, **p<0.01, ***p<0.001, ***p<0.0001, ns = non-significant.

**Figure 5:**
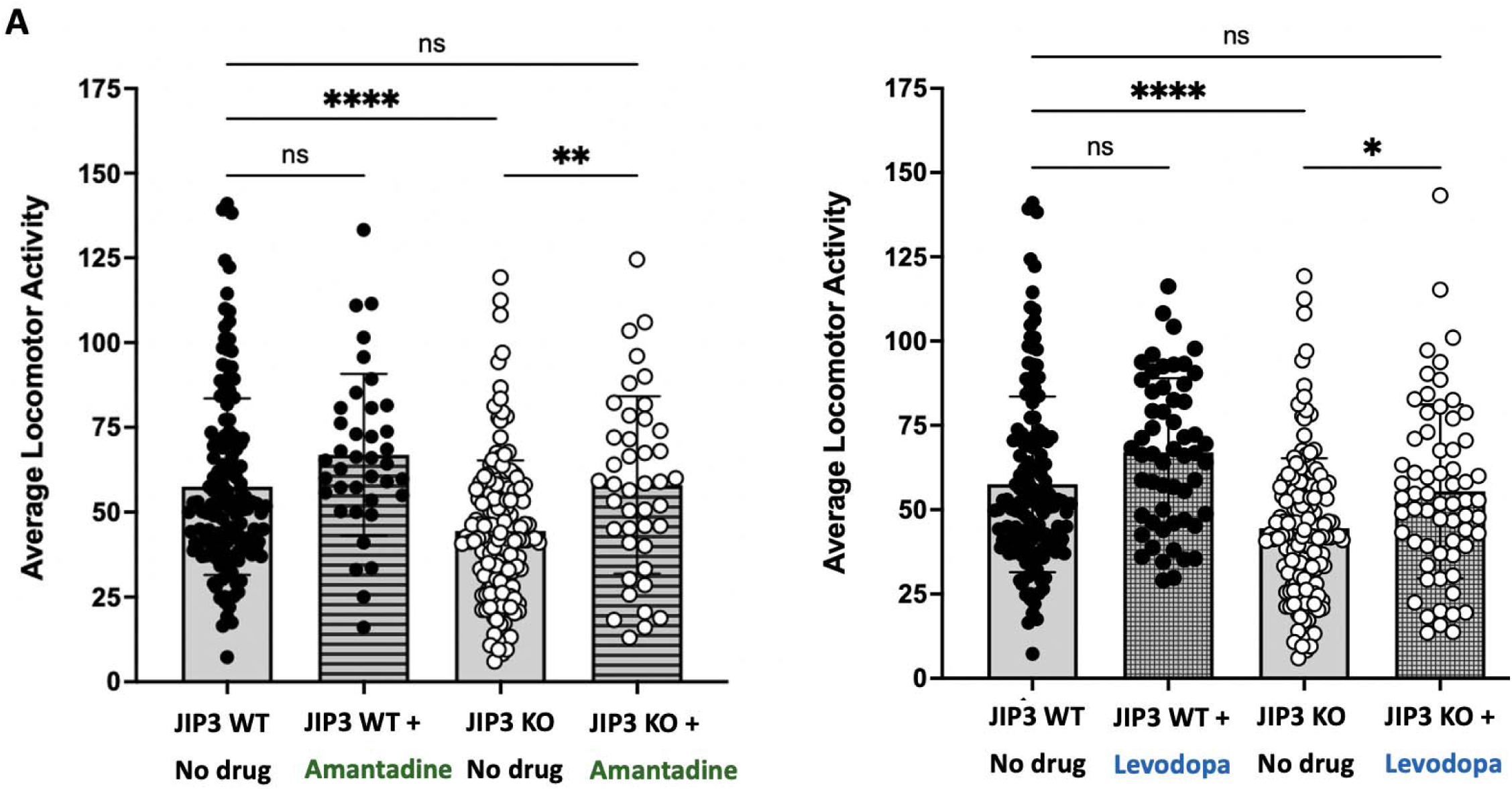
JIP3 KO locomotor phenotype rescued with anti-Parkinsonian drugs amantadine and levodopa. 2-hour MicroTracker assay displaying rescue of decreased locomotor activity in 6 dpf zebrafish larvae treated with 25 µM amantadine (A) and 100 µM levodopa (B). Each dot represents an individual zebrafish with mean and standard deviation provided. A 1-way ANOVA with Tukey’s test for multiple comparisons was conducted, p-values are *p<0.05, **p<0.01, ***p<0.001, and values > 0.05 were labeled as ns = non-significant.

We next began screening FDA-approved small molecules hypothesized to alleviate patient symptoms or target the disorder at the molecular level. These experiments showed that administration of amantadine or levodopa, immediately after fertilization, rescued average locomotor activity in JIP3 knockout larvae zebrafish (**Figure 6**). Both drugs lead to a slight increase in average locomotor activity in wild-type fish, but not to a significantly significant extent.

**Figure 6:**
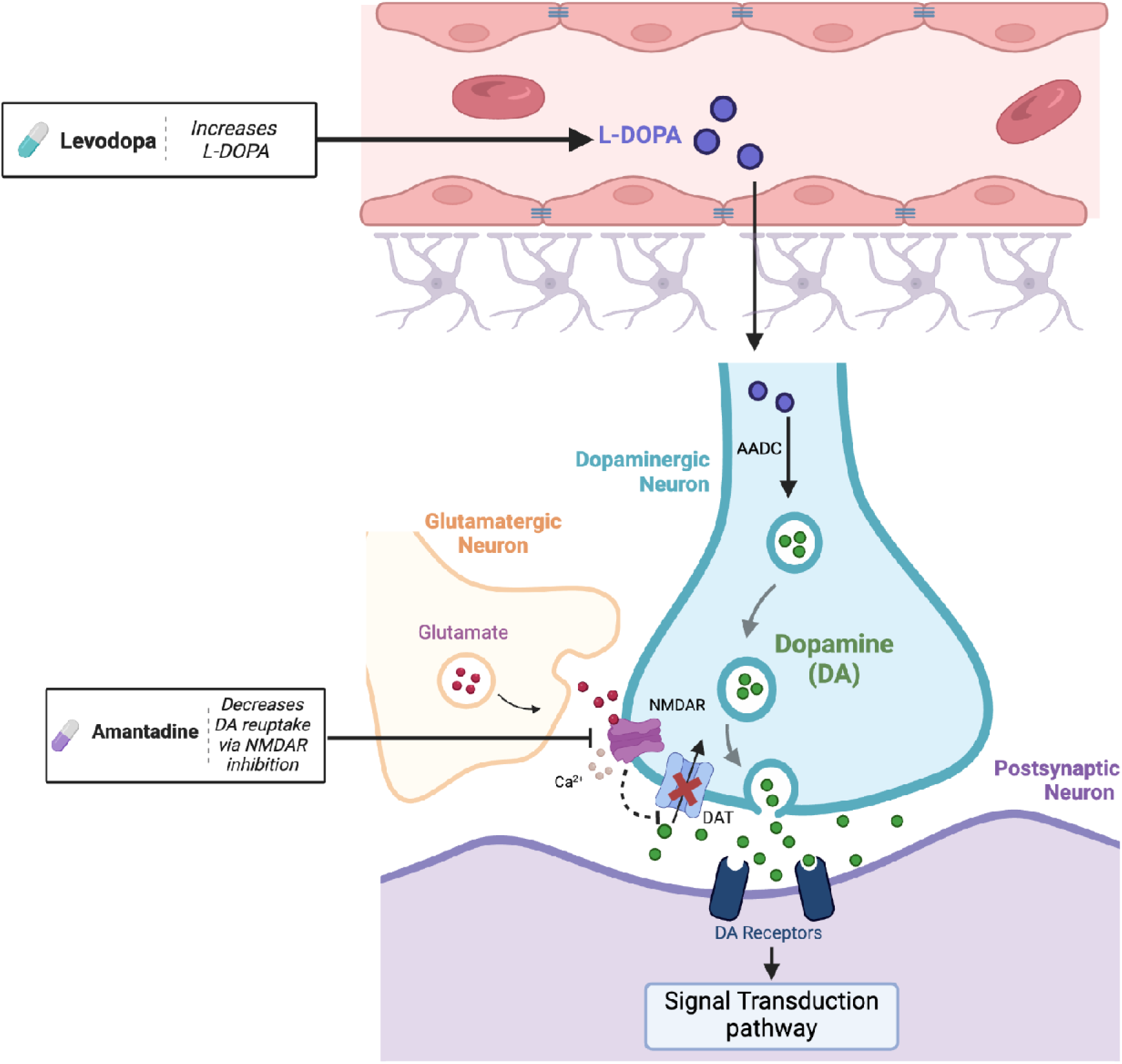
Treatment with levodopa and amantadine independently lead to an increase in dopamine in the synaptic cleft. Levodopa (L-DOPA) travels through the bloodstream and is transported across the blood-brain barrier, leading to increased levels of dopamine release. Indirectly, amantadine has been shown to decrease dopamine reuptake through inhibition of NMDA receptors which indirectly reduces dopamine transporter (DAT) activity, thus increasing dopamine levels at the synapse. We hypothesize that motility deficits are being rescued through these mechanisms of increased dopaminergic signaling. Created with Biorender.com.

## DISCUSSION

*MAPK8IP3*-related disorders, characterized by neurodevelopmental and motor impairments, are poorly understood at the molecular level, and no targeted treatments have been identified. In this study, we leverage an existing JIP3 KO zebrafish model to examine behavioral phenotypes and morphological abnormalities found in patients. We also establish a medium-throughput assay for screening of FDA-approved small molecules that are safe for use in children. These efforts demonstrate disease phenotypes in a JIP3 KO zebrafish model and provide preclinical evidence for candidate therapies for clinical trials in patients with *MAPK8IP3*-related neurodevelopmental disorders.

We identified gross morphological, behavioral, and locomotor deficits in JIP3 KO zebrafish throughout development. These deficits include reduced weight and standard length relative to wild-type fish and a potential sex-specific decrease in critical swimming speed. Importantly, these findings parallel clinical observations in patients with *de novo MAPK8IP3* variants, particularly reports of stunted growth, short stature, and motor abnormalities (Platzer et al., 2019). Potential sex-specific differences in motor function have not been reported in patients but warrant future study. Our overall findings are also consistent with a *Drosophila* model, where disrupted JNK signaling due to loss of JIP3 homolog *Syd* caused fly larvae to crawl significantly slower than controls, indicating impaired muscle function (Schulman et al., 2014).

Artificial intelligence and clinician recommendations identified levodopa and amantadine as candidate therapies to treat motor and behavioral abnormalities in these patients. These medications are effective and widely used to treat motor symptoms caused by dopaminergic deficits in patients with Parkinson’s disease. Here we found that levodopa and amantadine both rescued motor phenotypes in JIP3 KO zebrafish larvae, highlighting their potential for addressing motor dysfunction associated with the broader spectrum of *MAPK8IP3*-related disorders. One mechanistic hypothesis is that motor impairments from these disorders may be mitigated by enhancing dopaminergic neurotransmission during early development. We initially hypothesized that amantadine’s effects on locomotion phenotypes might be linked to its influence on BDNF levels, given that JIP3 facilitates BDNF transport, which regulates key downstream pathways (Huang et al., 2011; Ma et al., 2017). However, recent evidence suggests that BDNF loss enhances motility in zebrafish, indicating that other pathways may be involved (Lucon-Xiccato et al., 2023). While our results provide preliminary evidence for the therapeutic potential of these drugs, further research is needed to define critical windows for intervention and the mechanisms underlying the rescue of behavioral phenotypes.

Diverse mechanisms may underlie the pathogenesis of specific patient variants. Early truncating variants, such as p.(Y37X), are predicted to yield JIP3 loss of function, but additional experiments are needed. Furthermore, most of the reported *MAPK8IP3* variants are missense, and their functional consequences remain unclear. Future studies should investigate the possibilities of gain-of-function or dominant-negative effects, as well. Future work should also query potential deficits in dopaminergic circuitry in affected patients with Dopamine Transporter (DaT) scans.

In summary, we provide an in-depth characterization of morphological, locomotor, and behavioral deficits in JIP3 KO zebrafish and use medium-throughput drug screening to identify candidate therapies with implications for children with *MAPK8IP3*-related neurodevelopmental disorders. Future mechanistic studies are needed to examine how loss of JIP3 function alters neurodevelopment at the molecular level. Prospective clinical studies are required to formally evaluate the potential utility of levodopa and amantadine in patients with *MAPK8IP3*- related neurodevelopmental disorders or JIP3 loss of function conditions.

## ACKNOWLEDGMENTS

We thank Dr. Alex Nechiporuk and his team at Oregon Health and Sciences University for providing us with their JIP3 KO zebrafish line used in all analyses. We are grateful for the clinical guidance provided by Dr. Caroline Martinez at Mount Sinai School of Medicine and Dr. Harrison Walker at the University of Alabama at Birmingham Heersink School of Medicine. We would also like to thank Drs. Brad Yoder, CJ Haycraft, and Laura Lambert for their contributions to the scientific rigor and hypothesis generation for JIP3 loss of function. We are grateful to all the research teams associated with the Wolverine Foundation for their constructive feedback on experiments and data analyses.

## AUTHOR CONTRIBUTIONS

CC and PS designed experiments. CC, PS, GS, KT, SR, AA, RB, JB, AL, and AG performed experiments in zebrafish. CC, AF, AC, and MM assisted with AI-assisted drug candidate lists and analyses. CC, AF, PS, GS, KT and SR prepared figures. AF, CC, PS, GS, KT, and AL prepared the initial draft of the manuscript with help from AL. CC and MM secured funding for the project. CC directed the project and led the efforts to prepare the manuscript.

